# The Endoplasmic Reticulum as an Active Liquid Network

**DOI:** 10.1101/2024.05.15.594381

**Authors:** Zubenelgenubi C. Scott, Samuel B. Steen, Greg Huber, Laura M. Westrate, Elena F. Koslover

## Abstract

The peripheral endoplasmic reticulum (ER) forms a dense, interconnected, and constantly evolving network of membrane-bound tubules in eukaryotic cells. While individual structural elements and the morphogens that stabilize them have been described, a quantitative understanding of the dynamic large-scale network topology remains elusive. We develop a physical model of the ER as an active liquid network, governed by a balance of tension-driven shrinking and new tubule growth. This minimalist model gives rise to steady-state network structures with density and rearrangement timescales predicted from the junction mobility and tubule spawning rate. Several parameter-independent geometric features of the liquid network model are shown to be representative of ER architecture in live mammalian cells. The liquid network model connects the time-scales of distinct dynamic features such as ring closure and new tubule growth in the ER. Furthermore, it demonstrates how the steady-state network morphology on a cellular scale arises from the balance of microscopic dynamic rearrangements.

**SIGNIFICANCE:** The peripheral endoplasmic reticulum (ER) forms a continuous, dynamic network of tubules that plays an important role in protein sorting, export, and quality control, as well as cellular signaling and stress response. Elucidating how the unique morphology of the ER arises and supports its function is critical to developing a mechanistic understanding of the many neurological diseases associated with ER structural perturbations. We develop a physical model of the ER as an active liquid network to understand how its cellular-scale structure emerges from small-scale dynamic rearrangements. The model demonstrates how key features of ER architecture can arise from a balance of tubule growth and tension-driven sliding. This work provides insight into the fundamental physical mechanisms underlying the emergent morphology of the ER.

## INTRODUCTION

The endoplasmic reticulum (ER) consists of a vast inter-connected web of membrane-bound tubules and sheets in eukaryotic cells. It forms connections with many subcellular structures [1, 2], synthesizes and delivers lipids to other organelles [3, 4], stores and releases calcium [5, 6], and serves as a hub for the translation, folding, and quality control of secreted proteins [7, 8]. The ER is highly dynamic and assumes a variety of structural motifs, which aid in accomplishing these diverse functional roles and maintaining its interconnection with other organelles [9].

Prior work on ER structure has focused on how its diverse morphologies (tubules, junctions, helicoidal ramps, fenestrated sheets, cisternae, etc. [10]) arise from an interplay of ER morphogen proteins and membrane mechanics. A variety of ER membrane proteins (including the reticulons and the DP1/REEP/Yop1p family) induce and stabilize the high positive curvature of tubules [11, 12]. Others (e.g.: Climp63) stabilize the thickness of sheets and tubules [13, 14], while proteins such as lunapark [15] and the atlastin GTPase family [15– 17] help to form and maintain junctions. It has been shown that both tubules and sheets can be generated by curvature-producing proteins with their relative abundance determined by the absolute concentration of these proteins [13]. In other work, it was found that a diverse set of ER morphologies can be generated by tuning the proportions of reticulons and lunapark [18]. Recently, a model was developed which highlighted the role of intrinsic membrane curvature and ultra-low tensions in generating ER tubular matrices, ER sheet nanoholes and other intricate membrane structures [19]. These studies have helped elucidate the diverse structures observed in the ER as manifestations of local mechanical equilibrium.

Other work has sought to understand the link between ER dynamics and its structure. For instance, the anomalous diffusion of ER exit sites (ERES) along tubules is captured by a model of an individual ER tubule as a semi-flexible polymer [20]. The fluctuations of individual tubules have also been quantified, with variations in their dynamic behavior associated with different regions of the cell [21]. The dynamics of the ER are also implicated in controlling other subcellular structures; for instance, the motion of junctions may help regulate the distribution of microtubules within the cell [22]. Prior modeling work on plant cell ER [23, 24] demonstrated how small networks between persistent points appear to minimize the length of intervening tubules. Although these studies focused on small regions of the ER with only a few tubules, quantification of such minimal networks enabled estimation of biophysical quantities such as local membrane tension and viscoelastic properties of the cytoplasm. Notably, the plant ER was treated not as a polymer chain with spring-like stretching or bending energies, but rather as a network with constant tension along effectively fluid tubules, an assumption which we also adopt in this work.

The tubular network of the peripheral ER (Fig. 1A) undergoes dynamic rearrangements (Fig. 1B, Supplemental Video 1) that include two frequently observed processes. First, there is the creation of new tubules, which branch from and remain connected to the existing network. Most commonly, tubule creation occurs when cytoplasmic dynein or kinesin-1 motors bind to the ER and walk along acetylated microtubules [25, 26]. Other mechanisms of new tubule growth include attachment of the ER to the dynamic plus-ends of microtubules, deemed TAC events [27], or to motile organelles such as trafficking endosomes [28], lysosomes [29] and mitochondria [30]. The second class of dynamic ER rearrangements arises from the inherent membrane tension in the lipid bilayer of the tubules. This tension induces junction sliding and neighbor rearrangements, akin to the T1 rearrangements observed in foams [31], leading to a net decrease in network edge length [32]. As a result, loops of tubules can shrink until they vanish, often referred to as loop or ring closure. Certain loop closure events play important functional roles in the fission of mitochondria [30] and endosomes [33], while other loop closures are rapid and do not appear to be associated with other organelles.

**FIG. 1.**
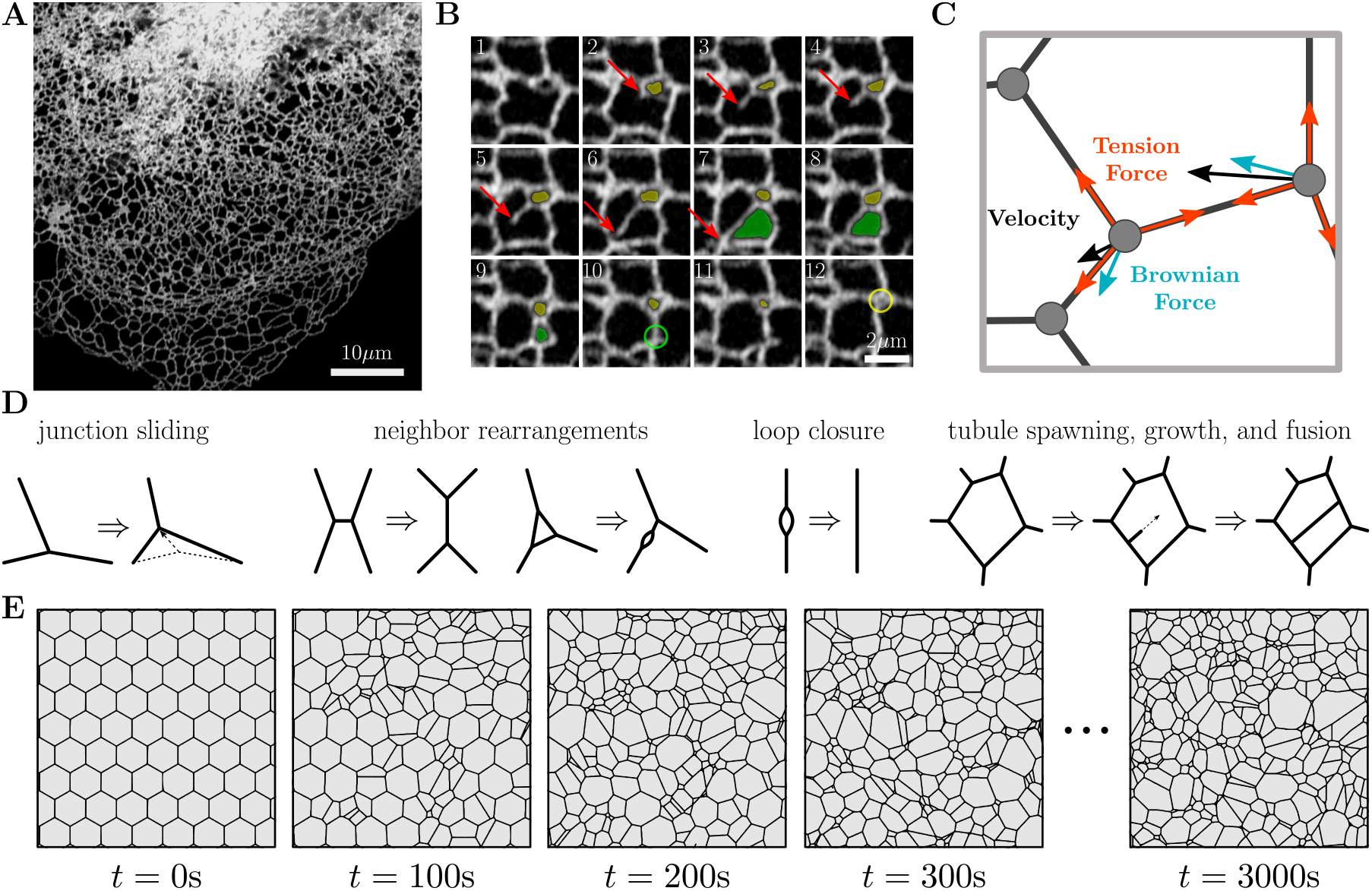
The dynamic structure of the endoplasmic reticulum is represented by a liquid network model. (A) Confocal image of COS7 cell expressing fluorescent endoplasmic reticulum (ER) marker (KDEL_mcherry), highlighting the peripheral ER network morphology. (B) Montage from same cell line illustrating the spawning of a new tubule (red arrow) and loop closure events (yellow and green shaded regions). Size of each image is 5.0 *×* 5.0 *μ*m with a time step of 0.6 s between each frame. (C) Physical model of junctions in a liquid network. A length-independent tension force (red) and a Brownian force (blue) combine to define junction velocities (black). (D) Possible rearrangements within a liquid network include junction sliding, neighbor swapping and rearrangement, loop closure and tubule spawning, growth, and fusion. (E) Representative simulated liquid network evolves from initially uniform honeycomb lattice to a random structure, with characteristic steady-state density. Region shown is a small segment of a much larger domain.

Similar structural rearrangements, such as T1 events and ring closures, have been observed in other physical systems characterized by interfaces under an effective tension. These include foams [34, 35], crystal grain growth in metals [36, 37], and patterns formed by mass-conserving reaction-diffusion systems [38, 39]. The evolution of domain boundaries in these systems is driven by a curvature-dependent effective pressure and a characteristic rate of material transport across the boundary. The tendency of large domains to grow at the expense of small ones leads to domain coarsening over time with a characteristic scaling [31, 40]. Other space-tiling systems, such as epithelial cell monolayers, do not coarsen, but also undergo boundary rearrangements driven by a combination of interface tension and domain pressure [41]. Such monolayers exhibit T1 junction rearrangements, ring closure due to cell extrusion, and domain splitting arising from cell division. Domain splitting is also observed in ER networks, resulting from the spawning and growth of new tubules. However, the ER differs from previous classic models of foams, crystal grains, and monolayers due to the ‘empty’ nature of the domains between tubules, with no substantial resistance expected to the exchange of cytoplasm between neighboring domains.

With the aim of connecting the small-scale dynamic rearrangements of the peripheral ER to its large-scale topological structure, we develop a simple physical model of the ER as an ‘active liquid network.’ We first identify junction mobility and tubule spawning rate as the primary regulators of steady-state network density within the liquid network model. The balance of creation and annihilation of tubules leads to a network structure with a characteristic density and connectivity profile. Key geometric quantities from the ER of living cells, such as the distribution of areas between tubules and their shapes, are reproduced by this simple model. Liquid networks with physiological densities are also found to rearrange at a rate consistent with the living ER. Intriguingly, extracted laws for growth and shrinking dynamics in simulations are sufficient to recapitulate the distribution of areas, thus highlighting how large-scale structure emerges from local dynamics. Cells may modulate these properties by tuning their effective junction mobility or tubule spawning rate. This could be achieved by static tethering of the ER to the cytoskeleton and other organelles and by withdrawal of newly growing tubules in ‘catastrophe’ events whose rate is quantified via semi-automated tracking in COS7 cells. Through computational models, analytic calculations, and quantitative image analysis of the peripheral ER in living mammalian cells, we identify physical rules governing its formation and maintenance on a cellular scale.

## RESULTS

### Emergent network topology from tension and growth

Inspired by the dynamic processes observed in living animal cells and prior descriptions of plant ER subregions as minimal networks [23, 24], we build a physical model of the large-scale mammalian peripheral ER as an active liquid network. The liquid network is composed of edges which transmit a membrane tension force between neighboring junctions. The membrane tension and tubule radii are assumed to be constant throughout the network.

In the low Reynolds-number environment of the cytoplasm, the motion of junction node *i*, located at position 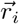, is assumed to obey an overdamped Langevin equation (Fig. 1C):

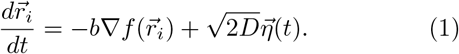

Here, *b* is the junction mobility (units of *μ*m/s), 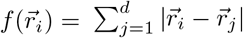 is the total tubule length connecting junction *i* with its neighbors at positions 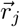, *D* is the junction diffusivity, and 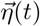 is a Gaussian distributed random noise term with zero mean and unit variance in each dimension. The first term describes the deterministic response to length-minimizing tension forces along the edges, and the second term describes a random Brownian force meant to capture the noisy environment of the cytoplasm.

With the above laws of motion, liquid network junctions are frequently pulled into close contact. Within some threshold distance of one another, junctions may swap neighbors, allowing for T1 transitions that lead to a decreasing edge length (details in Supplemental Material). Example neighbor rearrangements are shown in Fig. 1D. Occasionally, loops may form due to these neighbor swapping events, wherein two junctions are doubly-connected. These loops contract over time, due to there being twice as much tension pulling them together as there is pulling them apart.

In addition to the tension and diffusion driven motion of junctions, new tubule nucleation from existing tubules is modeled as an exponentially distributed random process with rate *k* (tubule spawning rate, units of *μ*m^*−*1^s^*−*1^). Newly spawned tubules grow with a velocity *v* (*μ*m/s), until coming into contact with an existing tubule, at which point the growth ceases and the tubules fuse, forming a stable junction.

In the endoplasmic reticulum, the velocity of growing tubules has been measured to be on the order of 1 *μ*m/s [26, 42]. We assume that the velocities of growing tubules are fast compared to the junction mobility (*v ≫ b*), so that a newly spawned tubule will span across a polygon before it substantially changes its shape. This assumption, justified via subsequently described measurements, allows us to treat tubule growth as nearly instantaneous. Additionally, the diffusive motion of junctions is assumed to be relatively small compared to experimentally observed tension-induced sliding events. With these assumptions, the behavior of the liquid network model can effectively be described by two parameters: the junction mobility, *b*, and the tubule spawning rate, *k*.

We simulate active liquid networks for a variety of parameter values, by integrating Eq. 1 forward in time (details in Methods). As seen in Fig. 1E, an initially uniform honeycomb network develops into a random network over time. The network continuously evolves, but after a transient initial period, the tubule density remains relatively unchanged. Thus, a steady state emerges due to the competing forces of junction motion controlled by edge tension and novel tubule spawning triggered by a random Poisson process.

### Steady-state density and timescales of liquid networks

Because the liquid network model is governed by only two parameters, dimensional analysis indicates there is a single characteristic length scale 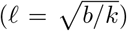 and time-scale 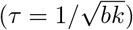 that define its behavior. Thus, changing the ratio of *b* and *k* generates networks with varying steady-state density and changing the product of *b* and *k* affects the dynamics of the network (Fig. 2A, Supplemental Video 2). To illustrate this effect, the results from nine simulations of liquid networks are shown in Fig. 2B. Each of the nine simulations has the same initial condition but a different combination of 𝓁 and *τ*.

**FIG. 2.**
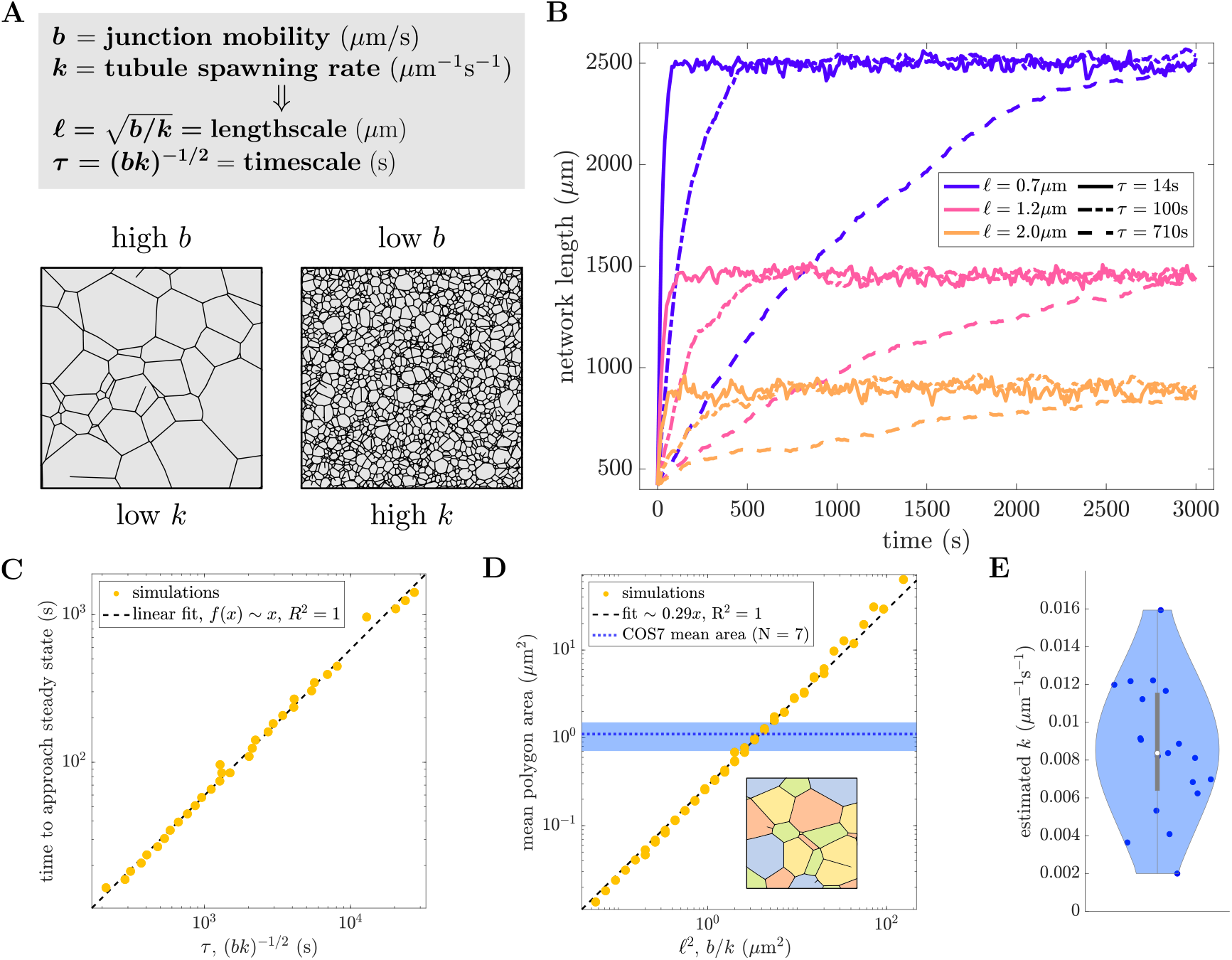
Steady-state network density and rearrangement timescales are set by two parameters. (A) Junction mobility and tubule spawning rate set the length and time scales (*𝓁, τ* respectively). (B) Network length over time for nine separate simulations, each starting from an initially sparse honeycomb network enclosed in a circle of radius 15 *μ*m. The steady-state network length is set by *𝓁* (color) and the time to reach steady state is set by *τ* (line style). Other parameters are *D* = 10^*−*5^ *μ*m^2^/s and *v* = 1*μ*m/s. (C) Timescale to approach steady state, extracted via exponential fit of curves as in (B) is proportional to the intrinsic network timescale *τ*. (D) The mean area of polygons in a liquid network is proportional to *𝓁*^2^ = *b/k*. Mean polygon area in COS7 cells is indicated by dotted blue line; shaded region gives inter-cell standard deviation (*N* = 7 cells). Inset: illustration of polygon extraction, different colored regions correspond to individual polygons. (E) Measured spawning rates from 19 COS7 cells, with mean and inter-cell standard deviation of 8.5 *±* 0.8 *×* 10^*−*3^ *μ*m^*−*1^s^*−*1^. Scatter data for (C), (D) is from simulations with mobility ranging between 0.001 *−* 0.1 *μ*m*/*s and spawning rate ranging between 0.0005 *−* 0.05 *μ*m^*−*1^s^*−*1^.

The total network length approaches a steady-state value over time, and can be fit to an exponential of the form *f* (*x*) = *c* (1 *−* exp (*t/τ*_steady_)). The parameter *τ*_steady_ gives an estimate of the time for liquid networks to approach steady state. This extracted time scales linearly with *τ* over a wide parameter range, as expected for simulations with a single dominant timescale (Fig. 2C).

To measure steady-state density one can consider the area of polygons (minimal loops) within the network. The mean area of these polygons scales linearly with 𝓁^2^ = *b/k* (Fig. 2D). It is therefore possible to construct networks with the same density as observed in living cells simply by tuning 𝓁. From a linear fit of the mean area across a range of *b* and *k* values we obtain ⟨*A⟩ ≈* 0.29*b/k*. The overall steady-state density of liquid networks can be captured by a simple equation for the total network length as a function of the new tubule spawning rate *k* and mobility *b*. For simplicity, we consider a network of hexagonal polygons. The total network length *L* can either grow via spawning of new tubules across the polygons, or decrease due to junction sliding:

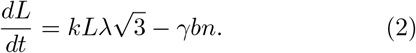

Here, *λ* is the average edge length, 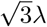 is the distance between parallel edges of a hexagon, *n* is the total number of junctions in the network, and *γb* describes the average sliding speed of a junction. At each individual junction, the value of *γ* must be between 0 and 3 depending on the angles between the adjacent tubules. The speed of growing tips is assumed to be sufficiently fast that growth events are instantaneous and there is no effect from the diffusion of junctions.

The total network length can be expressed in terms of the average edge length (*λ*) and the number of junctions (*n*) using the fact that the number of edges (*e*) in a hexagonal lattice is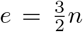. Thus, 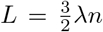 and at steady state 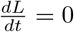 gives an average area of:

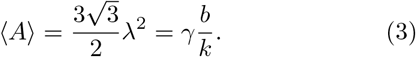

This predicted scaling of mean area with *b/k* aligns with expectations from dimensional analysis and agrees with the fit of simulation results.

The new tubule spawning rate *k* can be extracted directly from observations of ER dynamics in live COS7 cells (Fig. 2E, details in Methods). Together with the measured polygon area, this allows the effective mobility of the ER junctions in COS7 cells to be estimated as *b* = ⟨*A*⟩ *k*/γ = 0.03 *μ*m/s. We note that the relevant mobility is indeed much slower than the tubule growth speed (*b* « *v*), justifying our assumption of a single dominant length and timescale. One advantage of this approach is that tubule spawning rate is simple to directly measure from experimental data whereas junction mobility is a more opaque quantity. Thus, it is possible to calculate ER junction mobility via mean polygon area and spawning-rate measurements by taking advantage of the steady-state properties of liquid networks.

### Scale-invariant network structure reproduces ER morphology

While the two parameters of the liquid network model set length and time scales, any dimensionless metric of network structure must be parameter-independent. In particular, we consider the full distribution of polygon areas for a range of 𝓁 values (Fig. 3A). As the ratio of mobility to spawning rate increases, the distributions shift to the right, indicating an increase in large areas and a decrease in small areas. However, the overall shape of the distribution remains unchanged. At low areas, the log-binned distribution scales as 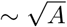 and at high areas it decays exponentially. When normalized by the mean area, the distributions collapse onto a single universal curve (Fig. 3A, inset), highlighting the scale-free nature of the model. Thus, increasing mobility is equivalent to “zooming in” on a patch of network, while increasing spawning rates leads to a denser network, or “zooming out.”

**FIG. 3.**
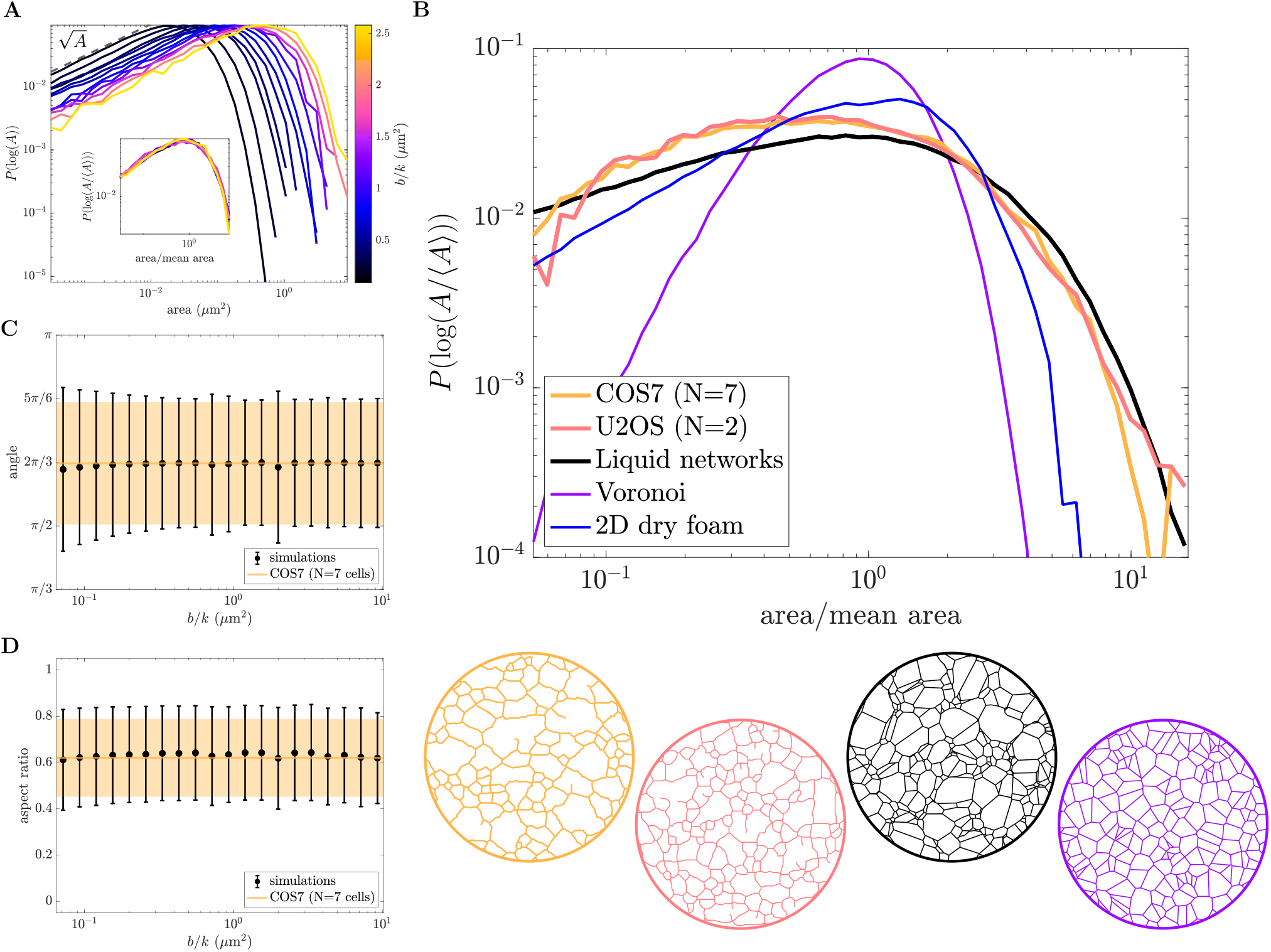
Scale-invariant liquid network model reproduces peripheral ER morphology. (A) Distribution of polygon areas in simulated liquid networks for a range of *b/k* values. Bin sizes are logarithmic. Inset: distributions collapse onto a single curve when normalized by mean area, revealing scale-invariant behavior of liquid networks. (B) Distribution of normalized polygon areas from the ER networks of COS7 and U2OS cells (yellow and pink, respectively) exhibit similar scaling to simulated liquid networks (black). For comparison, the distribution of areas for Voronoi networks and experimental measurements of a two-dimensional dry foam (data from [43]) are also shown (purple and blue lines). Example networks in matching color below. (C) Mean and standard deviation of angles between 3-way junctions in liquid networks (black) and the COS7 ER (yellow). (D) Mean and standard deviation of aspect ratio (shortest dimension/longest dimension) in liquid networks (black) and the COS7 ER (yellow).

Notably, the parameter-independent shape of the polygon area distribution for liquid networks provides a good approximation to that of the ER in living cells. In Fig. 3B, area distributions extracted from two different cell types commonly used to study the peripheral ER are shown (details in Methods). Both COS7 and U2OS cells (monkey kidney and human osteosarcoma, respectively) exhibit remarkably similar scaling, collapsing onto a single curve. The exponential drop-off at large areas is clearly conserved across experiments and simulations. The experimental measurements exhibit a slight enhancement of small-area polygons as compared to liquid network simulations. However, limitations in imaging resolution and segmentation prohibit the extraction of a reliable power-law for small-area scaling in the distribution.

The distributions for two other families of network are provided in order to demonstrate that the close match to ER morphology is not exhibited by other commonly studied 2D networks. The area distribution of simulated Voronoi networks (purple line) is comparatively narrow and sharply peaked around the mean. Experimental imaging data of a two-dimensional dry foam [43] (blue line), shows that the area distribution of foams is broader than Voronoi networks but still narrower than the ER and liquid networks. The foam also exhibits an exponential drop-off at large areas but has a linear scaling in the logarithmic distribution of small-area polygons.

It is possible to estimate the functional form of the liquid network area distribution (dashed black line) using only the rules for polygon growth and splitting, as described in the following section.

Other metrics of shape further confirm the similarities between the liquid network model and experimental ER networks. The distribution of angles between neighboring 3-way junctions is centered at 2*π/*3 or 120^*°*^. This is a universal property of two-dimensional networks composed of degree 3 junctions. The standard deviation of angles at 3-way junctions is also shown to be similar for simulated and ER networks across a wide range of parameter values (Fig. 3C). Finally, the aspect ratio of polygons (ratio of shortest dimension to longest dimension) is calculated in both simulated liquid networks and experimental ER networks. This provides another dimensionless, scale-free measure of shape: the mean and standard deviation of polygon aspect ratio are constant over a wide range of parameter values and approximately match experimental measurements (Fig. 3D).

### Model dynamics determine network rearrangement rates

Having demonstrated that the liquid network model approximately matches the steady-state morphology of ER networks, we next proceed to compare the dynamics of simulated and observed network structures. We consider two metrics for the dynamic rearrangement of the networks, demonstrating how parameters extracted from measuring polygon size and new tubule spawning enable accurate prediction of the time-evolution of ER network structure.

#### Edge mean minimal distance

To quantify the motion of individual tubules over time and the resulting changes in network structure, the edge mean minimal distance (EMMD) is calculated between the first and all subsequent networks, as described in Methods. Briefly, the network is meshed and for each point on subsequent networks, the minimal distance to a point on the starting network is found; these minimal distances are averaged to find the EMMD. The EMMD grows over time with an initial jump between the first and second frames that can be attributed to localization error and network segmentation artifacts (Fig. 4B, left panel). The same calculation is performed for simulated liquid networks with a matching average polygon size and a range of timescales.

**FIG. 4.**
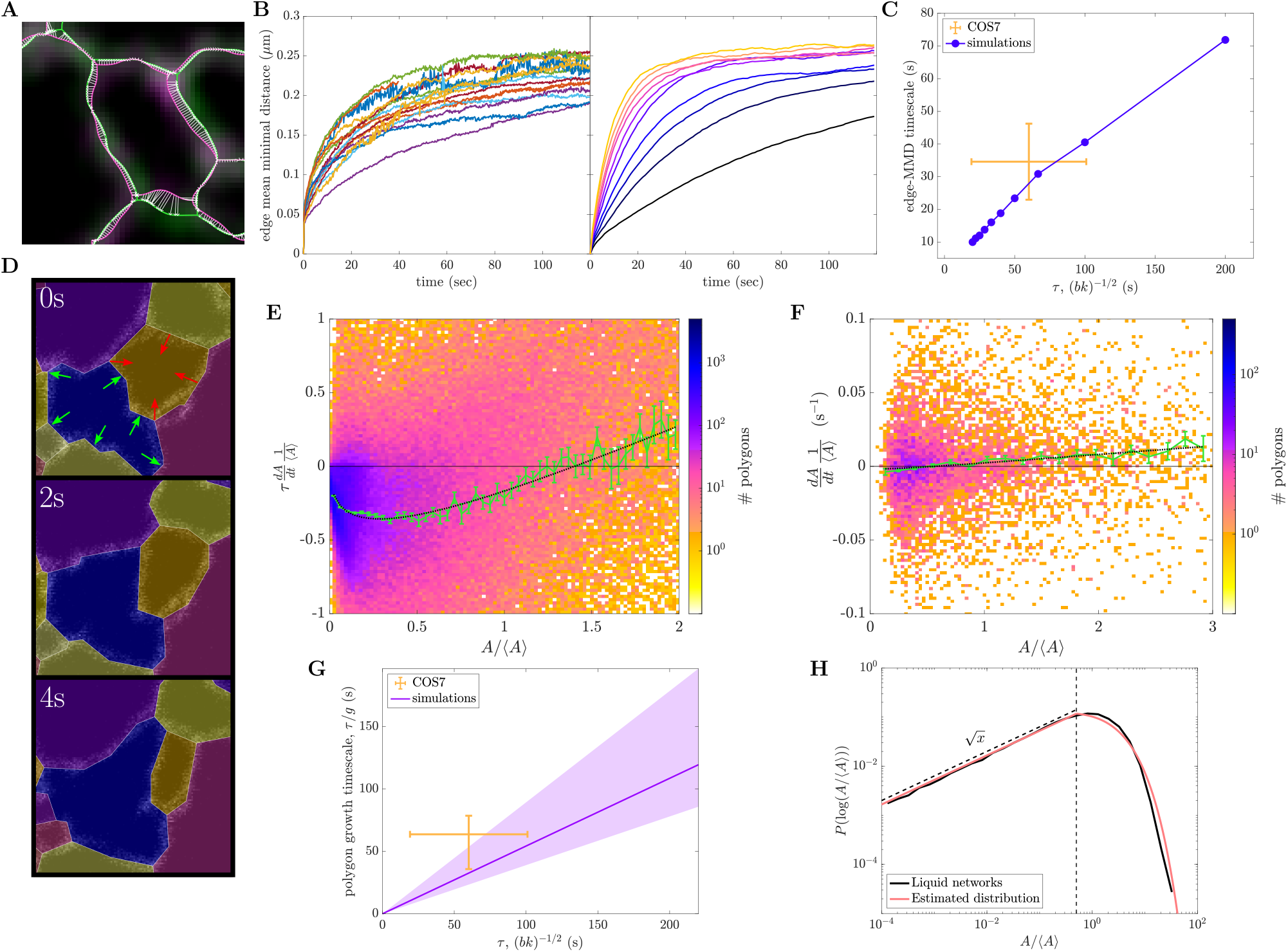
Dynamics of liquid networks predict ER rearrangement timescales and give rise to emergent polygon area distribution. (A) Confocal images of COS7 ER with 1s interval between first and last frames (green, pink). Extracted networks overlaid in matching colors. White arrows indicate minimal distance from meshed points on final network (pink) to meshed points on the initial network (green). (B) Left panel: edge mean minimal distance (EMMD) over time for 16 peripheral ER networks of COS7 cells. Right panel: EMMD of liquid networks with *τ* = (*bk*)^*−*1*/*2^ varying across one order of magnitude and fixed *b/k* = 4 *μ*m^2^. (C) Exponential timescale for EMMD to approach steady-state scales with *τ* for liquid network simulations. Yellow lines show mean value and intercellular standard deviation for COS7 cells. (D) Example polygon tracking of experimental data to quantify polygon growth and shrinking rates. Each frame is 5.0 *×* 5.0 *μ*m with a timestep of 2 s. Red and green arrows indicate shrinking and growing polygons, respectively. (E) Non-dimensionalized growth rates normalized by mean area and network timescale for 10 liquid network simulations. Green curve indicates mean (and standard error of the mean) within coarse bins of normalized area. Dotted black curve indicates fit to Eq. 4, giving *β* = 0.78, *g* = 1.69, *h* = 1.85. (F) Mean growth rates normalized by mean area and their dependence on relative polygon size from 7 experimental COS7 peripheral ER networks. Green curve indicates mean (and standard error of the mean). Dotted black curve indicates fit to Eq. 4, with *β* fixed at 0.78 giving *g/τ* = 0.018 s^*−*1^, *h/τ* = 0.015 s^*−*1^. (G) Polygon growth timescale (defined as *τ/g*) from 7 COS7 cells agrees with the simulated timescales extracted from simulations. (H) Distribution of normalized polygon areas in a simulated liquid network (black) compared to analytically approximated area distribution (pink). Vertical dashed line shows *A*^*\**^, where small-area and large-area solutions are joined.

Exponential fits of the EMMD growth over time give an effective timescale for the rearrangement of the network. As expected, this rearrangement time scales linearly with the intrinsic timescale *τ* of the simulations (Fig. 4C). The observed rearrangement timescale for the COS7 ER is 35 *±* 11 s, implying that during this time edges are significantly displaced from their original location. This timescale is of similar magnitude to the time required for the ER as a whole to explore a large fraction of the cytoplasm [44]. Notably, this rearrangement timescale is consistent with the simulation model, given the parameters *b* and *k* extracted from measurements of polygon area and tubule spawning rate. Thus, the liquid network model makes it possible to connect two seemingly independent dynamic processes—new tubule growth and rearrangement of existing edges.

#### Polygon growth rates

An additional metric to quantify the dynamic behavior of liquid networks arises from considering the growth and shrinking rates of polygons (independent of splitting events due to new tubule growth). Polygons are tracked in time and space using conventional particle tracking software [45] (details in Methods) We calculate the normalized growth rate for all tracked polygons in several COS7 ER networks and simulated liquid networks. The normalized growth rate is defined to be the time-derivative of each polygon’s area scaled by the average polygon area in that cell. For simulated networks, the growth rate is rescaled by the characteristic model timescale *τ* = (*bk*)^−1/2^, allowing simulations with different parameter values to be analyzed together. By determining the relationship between growth rate and polygon area the underlying laws governing the dynamics of the network can be probed.

A characteristic behavior of liquid networks is the growth of larger polygons and the shrinking of smaller ones (Fig. 4E). This is similar to the behavior of foams, where polygons with more than 6 sides grow while those with fewer sides shrink [34]. In liquid networks, as in foams [31] and crystal grain boundaries [37], larger polygons tend to have more sides and therefore to have internal angles above 120^*°*^, causing them to grow on average. As noted in Supplemental Material, the growth and shrinkage rates for a regular n-gon are expected to be <u>p</u> roportional to its perimeter, which scales roughly as 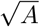. For large polygons, the number of neighbors should also increase as the polygon grows, giving a steeper dependence of the growth rate on area. We fit the average simulated growth rates to the following functional form (Fig. 4E):

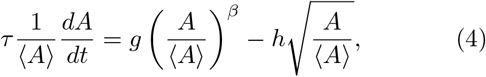

where the prefactor *g* encapsulates the rate of large polygon growth and *h* describes the rate of small-polygon shrinkage. From this fitted function, we can extract the typical rate *k*_grow_ = *g/τ* for the growth of large polygons.

When tracking polygons in images of live ER, the average growth rate is also negative for small polygons and positive for large ones, as in the liquid network model. Due to the limited data at small polygon areas, we fix the value *β* = 0.78 as fitted for simulated networks, and fit the remaining coefficients in Eq. 4 to the experimental data (Fig. 4F). This enables the extraction of a growth rate for large polygons, in real time units. As shown in Fig. 4G, the estimated growth rate for COS7 ER falls within range of the predicted value for liquid networks with the appropriate timescale *τ*. Thus, by measuring the average polygon area and rate of new tubule spawning (to set parameters *b, k*), the liquid network model makes it possible to predict the typical growth rate of large polygons in the ER network, thereby connecting distinct dynamic processes.

These results demonstrate that liquid networks are not only able to replicate key steady-state structural features but also capture the rearrangement timescales of living ER networks, providing a connection between morphology and tubule dynamics.

### Steady-state structure emerges from polygon dynamics

The dynamic behavior of network polygons can be abstracted still further by considering them as a population of individual aspatial entities capable of growing, shrinking, and splitting (Supplemental Video 3; details in Supplemental Material). The drift velocity of polygon areas *v*(*A*) = *dA/dt* is set by Eq. 4. The rate of splitting is expected to be proportional to the perimeter of a polygon, <u>w</u>hich scales as the square root of the area: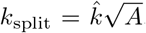. We define the distribution of polygon areas *P* (*A*), whose steady-state form must encompass a balance between small polygons disappearing due to shrinking and large polygons splitting into new ones.

In the limit of very small areas, the formation and disappearance of polygons due to splitting is negligible compared to the flux associated with polygon shrinking (see Supplemental Material for details). The overall rate of polygons disappearing when their area shrinks to zero can be estimated as:

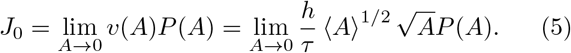

Because this flux must be a finite non-zero value, the small-area limit of the distribution is set to 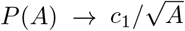, where *c*_1_ is a constant. A logarithmic transform gives the scaling *P* (log *A*)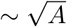, as observed in Fig. 3A.

In the limit of very large polygons, the distribution evolves due to area growth and splitting, and the steady-state form can be found by solving the resulting equation:

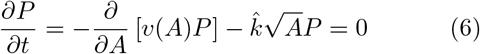

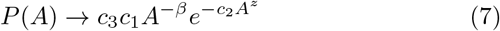

where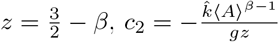, and *c*_3_ is a constant. In order to construct a full approximate distribution, the two limits for small and large area are married together at some intermediate value *A*^*\**^, thereby enforcing a value for the coefficient *c*_3_ = (*A*^*\**^)^1*−z*^ exp (*c*_2_*A*^*\*z*^). The coefficient *c*_1_ is obtained by normalizing *P* (*A*). Furthermore, the total rate at which polygons disappear (*J*_0_) and the rate at which new ones are produced 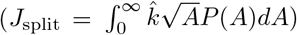 must be equal at steady state. This constraint fixes the transition value *A*^*\**^ *≈* 0.12*b/k*.

These calculations lead to a predicted average polygon area ⟨*A*⟩ ≈ 0.23(*b/k*), similar to the fitted relationship in Fig. 2D. Furthermore, the overall distribution of polygon areas, with a polynomial scaling at small A and an exponential scaling at large A, approximately reproduces the observed distribution from liquid network simulations (Fig. 4H). We note that the only fitting parameters employed in this analysis are the values of *g, h, β* in the expression for polygon growth rates as a function of area (Eq. 4). Thus measurement of polygon dynamics can be leveraged to approximately predict the steady-state architecture of the liquid network as well as the ER network structure in live cells.

### Pinning to static structures increases network density

We have shown how the structure and dynamics of liquid networks are governed by two parameters, the junction mobility and tubule spawning rate. In the next two sections, we explore how the cell effectively controls these parameters to modulate ER properties, first examining how junction mobility can be tuned by tethering of the ER to static structures.

The ER exists within the crowded, complex environment of the cytoplasm. It is pinned to the cytoskeleton via contacts with microtubules and actin filaments [46, 47]. The ER also forms critical contact sites with mitochondria [26, 30, 48], the plasma membrane [49], the Golgi [50, 51], endosomes [28, 33] and other organelles [1]. Quantification of ER network dynamics in plant cells has indicated that certain points along the network remain persistent over minute-long timescales [23]. To determine the effect of connections to static structures, we introduce a process for temporarily immobilizing junctions in liquid networks via pinning (rate *k*_*p*_). An unpinning process (rate *k*_*u*_) allows for a steady-state number of pins (Supplemental Video 4). The ratio *n*_*p*_ = *k*_*p*_*/k*_*u*_ sets the average number of pins in the model. We analyze systems with variable pin densities and with a wide range of pin persistence times.

To begin, we find how mean area depends on both junction mobility and pin density, calculated as *n*_*p*_ divided by total simulation area (Fig. 5A). For a fixed mobility, increasing the density of pins leads to a denser network with smaller mean areas. This effect is most pronounced for larger mobilities, leading to a steep decrease in mean area as pin density increases. The contour corresponding to the average area across COS7 cells is shown in green, indicating a wide range of possible mobility and pin density combinations in experimental networks.

**FIG. 5.**
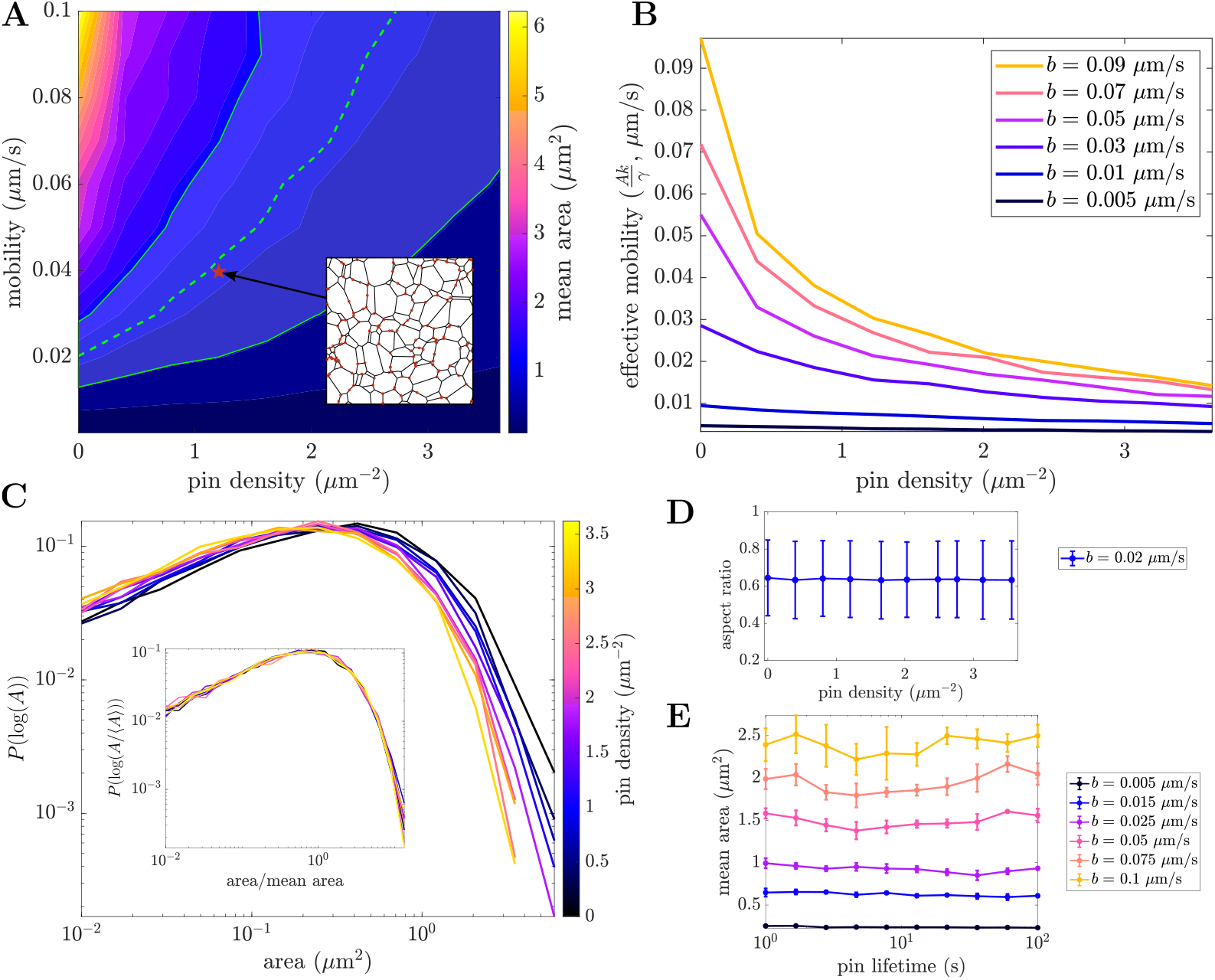
Tethering to static structures reduces effective mobility and increases network density. (A) Dependence of mean area on mobility and pin density. Dashed green line indicates contour corresponding to mean COS7 area, with shaded green region indicating standard deviation across all cells (N=7). Inset depicts an example 10 *×* 10 *μ*m snapshot of a liquid network with pinned nodes highlighted in red. (B) Effective mobility, defined as *b*_eff_ = ⟨*A*⟩*k*/*γ*, as a function of pin density for different mobilities. (C) Reversible pinning leads to denser networks, shifting the area distributions. Inset: normalized area distributions collapse to a single curve for all pin densities. (D) Aspect ratio is unchanged across a range of pin densities. (E) Mean area remains constant across a wide range of pin lifetimes. All results are with a tubule spawning rate of *k* = 0.005 *μ*m^*−*1^s^*−*1^.

The overall effect of pinning is to reduce the rate at which the network can relax and rearrange. For any given pin density, we can define an ‘effective mobility’ (*b*_eff_ = ⟨*A*⟩ *k/γ*, with *γ* = 0.29) by finding the value of *b* that would give the same average polygon area in the absence of pinned points. At low pin density, *b*_eff_ ≈ *b*, but as the pin density grows, effective mobility steeply decreases. In liquid networks, pinning to static structures or other organelles may thus limit network mobility and increase the corresponding density of ER network tubules in critical regions of the cell. For instance, this mechanism could aid the coalescence of ER around mitochondria contact sites.

There is little effect from pinning beyond tuning the density of the network. Area distributions shift as a function of pin density, but when normalized by mean area, the data collapse onto a single curve (Fig. 5C) just as in the case without pinning (Fig. 3A). There is also no effect on the mean and variance of polygon shape, as measured by aspect ratio (Fig. 5D). Furthermore, altering the pinning and unpinning rates (thus probing a wide range of pin lifetimes while maintaining a fixed pin density) has no effect on the steady-state structure of liquid networks (Fig. 5E).

### Tracking tubule spawning in COS7 ER reveals rate of catastrophe

Another mechanism through which cells can modulate ER network properties is by tuning tubule spawning rate. Within living cells, newly spawned tubules often cease growth and retract, a process we refer to as ‘catastrophe’. Independent of the underlying mechanism of growth (e.g. ER sliding or TAC events [25–27]), we quantify ER tubule catastrophe rates in COS7 cells, and explore the effect of catastrophe on network structure.

The tips of newly spawned tubules are tracked within COS7 cells (details in Methods). In addition to positional data, it is also recorded whether the growth is successful in fusing with a neighboring tubule (fused) or if it ultimately retracts (not fused). Using the position of tracked tips, the average velocity until fusion or until the time of retraction is calculated. The distribution of growth speeds is broad (Fig. 6A), with an average and standard deviation of 1.09 *±* 0.75 *μ*m/s, consistent with previous measurements [26, 42]. No significant differences in the distribution are observed between tubules that successfully fuse and those that do not.

**FIG. 6.**
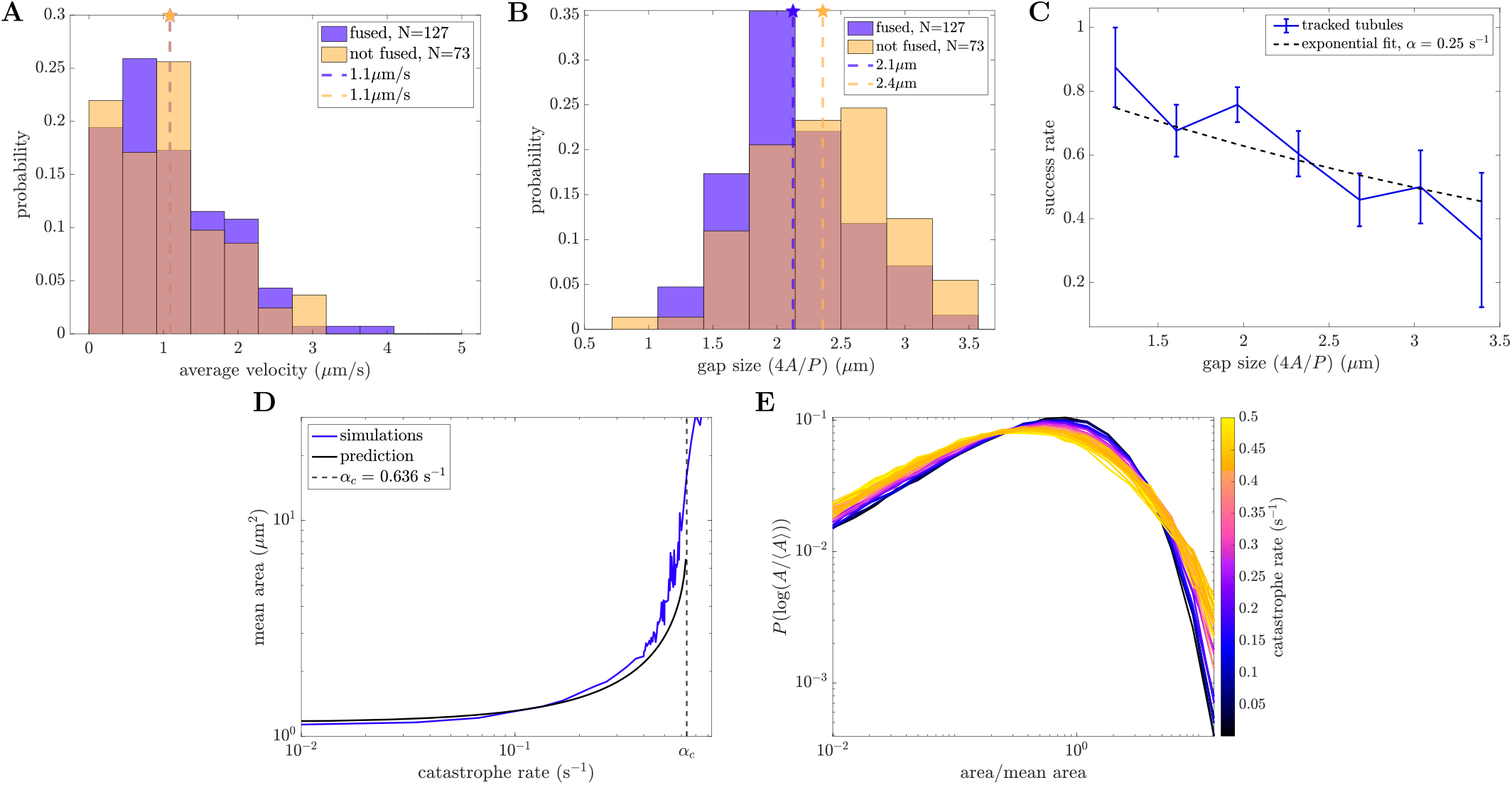
Growing tubules undergo catastrophe events that alter polygon size distributions. (A) Velocity distributions of growing tubules that lead to both successful and unsuccessful fusion. Mean of each distribution indicated by dashed line. (B) Distribution of gap sizes across which tubules are growing for both successful and unsuccessful fusion events. Mean of each distribution indicated by dashed line. *p* = 0.001 for a one-sided Student’s t-test. (C) Success rate as a function of gap size *d*, with fit to *S*(*d*) = *e*^*−αd/v*^ indicated by dashed line. (D) Mean area grows as a function of catastrophe rate. Black line marks analytic prediction. Dashed vertical line shows predicted critical rate beyond which mean areas diverge. (E) Normalized area distributions for liquid networks with increasing catastrophe rates. High catastrophe rate leads to an enrichment of both small and large areas.

We next calculate the gap size that growing tubules must traverse in order to fuse, a quantity which differs between successful and unsuccessful spawning events (Fig. 6B). Here, gap size is the approximate diameter of the enclosing polygon (*d* = 4*A/P*, with *A* area and *P* perimeter, an exact formula for the inscribed diameter of regular polygons). On average, successful growth events traverse shorter gap sizes (2.1 *±* 0.5 *μ*m) than unsuccessful events (2.4 *±* 0.5 *μ*m). This effect can also be visualized by considering the success rate of newly spawned tubules, which decreases as a function of gap size (Fig. 6C).

An estimate for the catastrophe rate in living ER networks can be extracted by fitting the experimental success rate to *S*(*d*) = *e*^*−αd/v*^ (Fig. 6C). Here, *d* is the gap size, *v* = 1.1 *μ*m/s is the average velocity of all measured growth events, and *α* is the catastrophe rate and fit parameter. This gives *α*_ER_ = 0.25 *±* 0.04 s^*−*1^.

To explore how these events affect the steady-state structure of the network, we introduce a constant-rate (Poissonian) catastrophe process for each growing tubule in the liquid network simulations. A single parameter (*α*) controls the rate at which a growing tip ceases forward motion and begins to retract due to membrane tension. Simulations are performed with experimentally relevant choices for mobility and spawning rate (*b* = 0.02 *μ*m/s, *k* = 0.005 *μ*m^*−*1^s^*−*1^) across a wide range of catastrophe rates, and the steady-state structural properties of the network are analyzed.

At small catastrophe rates, the mean area of polygons in the network remains relatively unchanged from the case with no catastrophe (Fig. 6D). As the catastrophe rate increases, it becomes more likely that growing tubules will retract before fusing, especially across larger gaps. Thus, there are fewer splitting events of large areas, leading to an increase in mean area. The larger average gap size, in turn, leads to even fewer fusion events. As the catastrophe rate grows sufficiently large, the persistence length of growth events becomes smaller than the average gap they must traverse. Mean area is then expected to diverge beyond a critical catastrophe rate, as confirmed by simulations (Fig. 6D).

The average effect of catastrophe on steady-state structure in liquid networks can be analytically approximated by modifying Eq. 2 for the total network length to incorporate the success rate *S*(*d*) of new growth events:

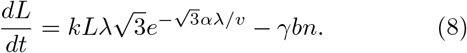

At steady state, this reduces to the following transcendental equation for *λ*,

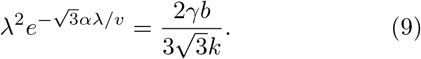

The derivative of *λ* with respect to *α* approaches infinity at 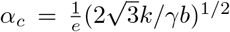, indicating that no finite solution is possible above this critical catastrophe rate. For the simulation parameters used in Fig. 6D, the critical value is given by *α*_*c*_ = 0.64 s^*−*1^. Solving for *λ* numerically, the polygon mean area can be found as a function of catastrophe (black curve in Fig. 6D). The divergence of average area with increasing catastrophe rate is successfully predicted by this analytic model.

Notably, the catastrophe process affects not only the average polygon area, but also the normalized distribution of areas (Fig. 6E). The altered distributions arise because catastrophes have a greater effect on large than on small polygons. Thus, normalized distributions for systems with frequent catastrophes have a fatter tail of large-area polygons that are unlikely to be split by a successful new tube fusion.

## DISCUSSION

In this work, we use a physical model of the peripheral ER as an active liquid network to capture its steady-state structural and dynamic properties. The behavior of liquid networks is effectively described by two main parameters, the junction mobility and tubule spawning rate. A characteristic network density and connectivity emerges from a balance between tubule creation and the contraction of small polygons. This model reproduces key geometric features of the peripheral ER in adherent mammalian cells, such as the typical shape and distribution of areas between tubules. We find that liquid networks are able to replicate physiological rearrangement timescales. Quantifying polygon dynamics in these systems allows us to derive the distribution of areas, thus forming a connection between emergent steady-state structure and the underlying, small-scale dynamics. Finally, by considering the effects of static tethering points and catastrophe of tubule growth we explore how the cell can alter the effective junction mobility and tubule spawning rate to modulate network properties.

In this work, the complex protein-studded membrane structure of the peripheral ER is reduced to a spatial graph of junctions connected by one-dimensional, constant-tension, fluid-like edges. This highly simplified model is able to recapitulate much of the structural and dynamic properties of the living ER, while remaining agnostic to the specific details of membrane-shaping and dynamics at the nanometer scale. In reality, the effective tension driving the shortening of ER tubules could be modulated by varying tubule radii and by the distribution of tubule-stabilizing proteins such as the reticulons [52]. Both tubule radii and reticulon distributions are spatially heterogeneous across the network [14], potentially giving rise to tension gradients. More complicated ER structures, such as fenestrated sheets [53], may also influence the tension and hence the dynamics of the surrounding network. Furthermore, tubule spawning rate may be heterogeneous due to varying distributions of microtubules and motors. These effects may account for some of the spatial variability in ER density observed in cells [54].

An additional simplifying assumption inherent to the liquid network model is the ability of individual tubules to straighten on a time-scale faster than the node rearrangements. While some bent tubules can be seen in the peripheral ER, their persistence length [21] is usually much longer than the typical polygon size. Thus the kinks that sometimes appear in long ER tubules are likely associated with connections to other cellular structures (analogous to the ‘pinning points’ discussed above). The constant, isotropic mobility coefficient for network nodes is another simplification of the model, neglecting the differences in friction associated with dragging a long tubule perpendicular to its axis, versus sliding a junction along the tubule membrane. Despite these simplifications, the liquid network model reproduces many aspects of ER structure and dynamics, helping to connect local rearrangements to the large-scale architecture of the network.

Maintaining an organelle as an active, dynamic liquid network incurs a continuous energy cost for the cell. In particular, motor activity or driven microtubule polymerization is required to grow new tubules that split polygons and interrupt coarsening. Given that individual kinesin motors burn one ATP molecule per 8-nm step [55], we would expect the rate of energy consumption associated with network maintenance to scale as 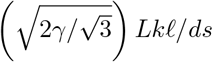, where *L* is the total network length, 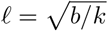 is the motor step size, and *γ* is the scaling factor relating average polygon area to 𝓁^2^. For the cells considered here this would amount to roughly 2 *×* 10^3^ ATP/sec. Notably, this maintenance cost is orders of magnitude lower than the estimated energy consumption (*∼* 10^8^ ATP/sec) associated with synthesizing the proteins shipped from the ER in the secretory pathway [56– 58].

For this modest energetic cost, the dynamic network of the ER provides the cell with a number of functional benefits. As a topologically isolated space with high calcium concentration, the ER provides a compartment for the efficient folding of proteins destined for the extracellular environment [7, 8]. Its dense network structure enables the rapid and proximal delivery of calcium ions into the cytoplasm [6] during localized signaling events known as puffs or sparks [59]. The well-connected network architecture also allows for rapid search by newly folded proteins to encounter exit sites in the ER [54]. Furthermore, the density of the network makes it possible for the ER to form a plethora of contact sites for the transport of proteins, lipids, and ions to and from other organelles such as mitochondria and endo/lysosomes [29, 30, 33, 60].

The dynamic rearrangements of the network could allow for rapid structural response of the ER to local and global perturbations. For example, the increased network density associated with pinning points may aid the accumulation of ER tubules near mitochondrial contact sites, where the ER is known to play an important role in mitochondrial fission and fusion [30, 61]. Network dynamics could also allow the ER to restructure around rearranging organelles or in response to cytoplasmic deformation in motile cells [62]. Finally, the liquid network would be able to reform rapidly following mitosis, during which the ER undergoes a global tubule-to-sheet conversion [63, 64].

The liquid network model not only accounts for the unique reticulated structure of the ER but also demonstrates how this architecture can emerge from and be regulated by a balance of two simple dynamical processes: tension-driven coarsening and new tubule growth.

## MATERIALS AND METHODS

### Liquid network simulations

To simulate liquid networks, the overdamped Langevin equation for network nodes (Eq. 1) is integrated forward in time using a fourth-order Runge-Kutta method implemented in Fortran 90.

After updating junction positions in each time step *dt*, the code checks for connected nodes which are within some minimal distance of each other, defined to be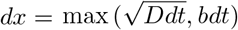, where *D* is the diffusivity and *b* the mobility of the nodes. This value tends to be on the order of 10^*−*4^ *−* 10^*−*3^ *μ*m. For connected nodes within this minimal distance, the length of all possible rearrangements of their neighbors is calculated, to determine which configuration minimizes the connected length (Fig. 1D). The minimal configuration is chosen, and edges accordingly undergo their small rearrangements.

Next, the code checks for fusion of actively growing tubule tips with existing edges by finding any crossings that may have occurred. Upon identification of a crossing event, the tubules fuse and form a stable junction node (Fig. 1D). Finally, the code samples from a Poisson distribution with rate *k* to determine the number of new tubule spawning events occurring in the current time step *dt*, given the current network length (excluding boundary edges). Novel spawning events occur uniformly distributed across all non-boundary edges of the network. Thus concludes one cycle of the simulation, after which the code iterates to update junction positions. Unless stated otherwise, a time step of *dt* = 0.01 s is used, the initial configuration is a honeycomb network, and the simulation is run until steady state has been reached (as defined by the stabilization of total network length over time).

To avoid runaway growth or full collapse of the membrane, all simulations are enclosed within a circle of radius 14 *μ*m defined by fixed immobile junctions connected via boundary edges. When a tubule growth event fuses with a boundary edge, the resulting boundary junction behaves exactly the same as junctions in the bulk except its motion is restricted to the perimeter of the boundary circle. Neighbor rearrangements and loop closure can still occur, resulting in a net balance of boundary junctions over time. The circular boundary is taken to represent the plasma membrane, which is known to make direct contacts with the peripheral ER network.

Extensions to the model include random static pinning and unpinning of bulk nodes, and the catastrophe process for tubule growth, all modeled as random Poisson distributed processes. Pinning of nodes occurs with rate *k*_*p*_ (s^*−*1^), and unpinning of nodes occurs with rate *k*_*u*_ (s^*−*1^). This pinning and unpinning is performed at the end of each simulation cycle. Catastrophe of each growing tubule occurs with rate *α* (s^*−*1^). Catastrophe events are sampled every cycle after checking for tubule fusion.

We obtain spatial network structures over time from these simulations in the form of node positions and edge connectivity. To analyze the properties of minimal loops or polygons in these network structures, we use publicly available code written in Matlab [65].

### Cell culture and imaging

COS7 cells (ATCC-CRL-1651) and U2OS cells (ATCC-HTB-96) were grown in Dulbecco’s modified Eagle’s medium (DMEM) supplemented with 10% fetal bovine serum (FBS) and 1% penicillin/streptomycin. Prior to imaging experiments, COS7 cells were seeded in six-well, plastic bottom dishes at a density of 150,000 cells per well, about 16 hours prior to transfection of DNA plasmids. The mCh-KDEL and KDEL-venus were described previously [66] and were transfected at a standard amount of 0.2 *μ*g for all experiments. Plasmid transfections were performed 24 hours prior to imaging with plasmid DNA in Opti-MEM (Invitrogen) with 5 *μ*L of Lipofectamine 3000 reagent (Invitrogen) according to the manufacturer’s instructions. After 5 hours of transfection, cells were seeded in 35 mm glass-bottom microscope dishes (MatTek).

All imaging experiments were performed at the Van Andel Institute Optical Microscopy Core on a Zeiss LSM 880, equipped with an Axio Observer 7 inverted microscope body, stage surround incubation, Airyscan detector, two liquid-cooled MA PMT confocal detectors, and one 32-channel GaAsP array confocal detector. Images were acquired with a Plan-Apochromat 63x (NA 1.4) oil objective with 3x optical zoom using Zeiss Zen 2.3 Black Edition software. Cells were tracked for at least 2 minutes with constant acquisition (200 ms or 316 ms).

### Image analysis

#### Extracting networks from live cell images

The machine learning segmentation toolkit ilastik [67] is used to segment ER network structures from live-cell images using the KDEL Venus or mCherry KDEL markers. A network structure is extracted from the probability file output by ilastik using custom code for skeleton tracing in Matlab [68]. This code is publicly available at https://github.com/lenafabr/networktools and includes data structures for storing the network as a set of nodes connected by edges with curved spatial paths.

#### Estimating the tubule spawning rate in living cells

All new tubule spawning events are manually counted within cropped subregions (6.3 *×* 6.3 *μ*m) of the peripheral ER from 19 cells over the course of 40 s (Supplemental Video 5). The number of tubule spawning events is then normalized by the average network length over all frames (as obtained via ilastik and Matlab skeletonization, see above) and the total time considered. This yields an estimate for the average tubule spawning rate, *k*, (units of *μ*m^*−*1^s^*−*1^) in each cell.

#### Extracting polygons from experimental ER images

The machine learning segmentation toolkit ilastik [67] is used to accurately identify polygons in experimental images of ER networks (7 COS7 cells and 2 U2OS cells). First, images are segmented as before using the pixel classification capabilities of ilastik. Then, an object classification model is trained to identify polygons in the cell periphery using the previously generated probability file and raw image as input. Associated geometric and spatial properties of these polygons are then analyzed in Matlab.

#### Semi-automated tracking of new tubule growth events

Network structures are extracted from images of COS7 cells, and all degree-1 (terminal) nodes are identified. The positions of these nodes and the length of their connected edges are calculated. The terminal nodes are then tracked over time using conventional particle tracking software [45] where the particle ‘amplitude’ is set by the connected edge length. Tracks within 1 *μ*m of the network boundary or with track length less than 3 frames are discarded. The tracks are manually verified and adjusted as necessary using a custom-built Matlab GUI that superimposes track data on top of image files. The fate of the growing tubule (whether or not it successfully fuses into the network) is manually identified. A total of 200 tubule tip trajectories are quantified for Fig. 6.

The average velocity is found by dividing the maximum distance a tubule travels from its spawning point by the time it takes to get there. For successfully fused tubules, this is simply the distance from spawning to fusion; for unsuccessful spawning events, it is the distance from spawning to the turnaround or catastrophe point. The area and perimeter of the polygon containing the spawning event are calculated for each tracked frame. The gap size is estimated as *d* = 4*A/P* and is averaged over the whole track.

### Quantifying network rearrangement

#### Edge mean minimal distance

Both simulated and experimental networks are meshed into 0.03 *μ*m segments, and for each point on subsequent networks, the minimal distance to a point on the starting network is found. The edge mean minimal distance (EMMD) is the mean of all such distances. To extract a timescale from the experimental data, the EMMD versus time curve is fit with a double exponential function of the form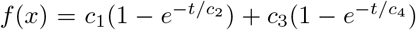. The longer timescale (e.g. *c*_2_ *≫ c*_4_) is chosen as the representative EMMD timescale for each experimental network. Simulated data does not exhibit the same short timescale jump as experimental because there are no localization errors or network segmentation artifacts. Thus, only a single exponential (e.g. the first term of *f* (*x*)) is fit to the simulated EMMD curves, giving the extracted EMMD timescale for each simulated network.

#### Polygon growth rates

Experimental polygon positions and areas are extracted from ilastik segmented images (see above). Matlab File Exchange code for extracting minimal loops from graphs is used to find polygons in simulated networks [65]. Polygons are tracked over time using conventional particle tracking software [45]. For each tracked polygon, we also perform a change point analysis of the tracked area over time using Matlab’s findchangepts function [68]. This enables identification of additional splitting events that may have been missed by the tracking software. The tracks are broken up at the change-points, allowing for robust tracking of polygon growth and shrinking dynamics between split events.

From these tracks, the area rate of change over 3 s intervals is calculated in both experiments and simulations after applying a lowess smoothing filter with span 3 s. Only tracks longer than 4s are analyzed. The normalized growth rate is obtained by dividing the extracted growth rate by the average area of all tracked polygons in that cell or simulation run. In order to collapse together simulations with different parameter values, we multiply the simulated growth rate data by the intrinsic network timescale *τ* = (*bk*)^*−*1*/*2^, yielding a non-dimensionalized growth rate.

The mean and standard error of the mean of normalized growth rate are calculated within bins of normalized polygon area (green points in Fig. 4E, F). This provides an estimate for the average growth and shrinking behavior for polygons as a function of their area within liquid networks and the ER. The binned mean growth rates are then fitted to Eq. 4. For simulation data, *β, g, h* are the fitting parameters, whereas for experimental data we fix *β* = 0.78 (the scaling exponent obtained from the simulation fits) and only fit *g, h*. In both cases, the data is resampled 2000 times to estimate the error in the fitting parameters, assuming normally distributed values with mean and standard deviation obtained from binned values.

## ACKNOWLEDGMENTS

We thank Massimo Vergassola for helpful discussions. We thank the Van Andel Institute Optical Imaging Core (RRID:SCR 021968), especially Lorna Cohen, for their assistance with the Zeiss LSM 880. Support for this work was provided by the National Science Foundation (Grant ID #2034482 to EFK and #2034486 to LMW), the Research Corporation for Science Advancement (EFK), Chan Zuckerberg Biohub San Francisco (GH), the Chan Zuckerberg Initiative Theory Hub (GH and EFK) and the Supporting Structures: Innovative Partnerships to Enhance Bench Science at CCCU Member Institutions program (SCIO, John Templeton Foundation, and the MJ Murdock Charitable Trust; LMW).

## SUPPLEMENTAL MATERIAL

### Polygon growth dynamics: regular n-gons

Informed by the equation of motion for junctions in a liquid network (Eq. 1), an approximate form for the growth law can be derived. For simplicity, we consider the growth dynamics of regular n-gons, whose outgoing connections are assumed to bisect the angle of each junction. The rate of change in area *A* is given by:

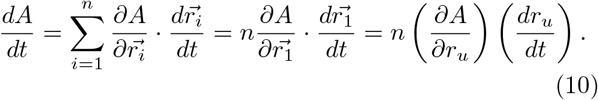

where 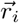 are the positions of each of the polygon’s *n* identical junctions, and *r*_*u*_ is the component along the radial direction.

Assuming diffusion is negligible, the junction motion is determined solely by the length minimization term from Eq. 1, giving

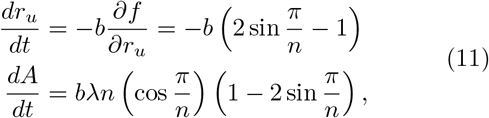

where *λ* is the side length of the polygon.

This predicts a negative growth rate for 3 *< n <* 6 and a positive growth rate for *n >* 6. For small polygons, we would expect *n ≈* 3, and the shrinking rate should scale with the polygon perimeter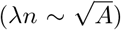. For large polygons, the rate of growth can be described as 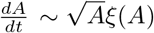 where *ξ*(*A*) grows with the number of polygon sides. Because the number of sides is expected to grow with area, *ξ*(*A*) should increase with *A*. We therefore make the approximation that large polygon growth rates increase as *A*^*β*^ for some approximate value of 0.5 *< β <* 1. Overall, we take an ansatz that linearly combines the two scaling relations for small and large area polygons:

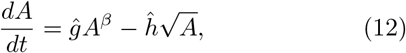

where normalization by the average area yields Eq. 4 in the main text. It should be noted that this analysis does not account for the irregular shape of the polygons in the liquid network and thus provides only an approximation for the growth rate as a function of area.

### Derivation of the polygon area distribution from growth and splitting laws

The goal here is to determine the steady-state distribution of polygon areas in a liquid network, *P* (*A*), starting from the laws for growth and splitting of individual polygons and the assumption that polygons on average take the form of a hexagon. The growth law extracted from simulations (Fig. 4F) and the splitting rate of these polygons can be expressed as follows:

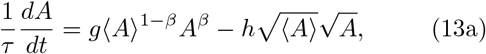

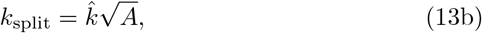

where *A* is polygon area, ⟨ *A ⟩* is the average area of all polygons, *g, h*, and *β* are constants determined from a fit of simulated growth rates, and 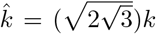 is the prefactor for splitting in terms of the spawning rate *k*. Equation 13b is obtained by finding the perimeter *p* of a regular hexagon of area *A* and noting that the expected splitting rate is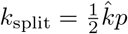, since new tubules can grow in either direction perpendicular to an edge.

If we make the additional assumption that polygons tend to split in half upon new tubule growth, the distribution of polygon areas, *P* (*A*), would be expected to obey an effective Fokker-Planck equation:

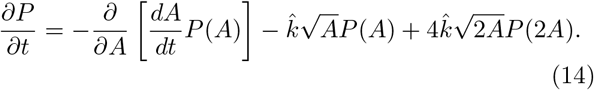

Here, the first term corresponds to the area drift associated with polygon growth and shrinking, the second term describes polygon disappearance due to splitting, and the last term corresponds to the formation of new polygons after a splitting event [69].

#### Small area scaling

Although Eq. 14 is not easily solvable analytically, we can analyze its behavior in the limit of very small polygons. We consider a narrow band of small polygon areas: (0, *δA*). The rate at which polygons disappear from this band due to shrinking below 0 area (*J*_0_) must equal the total rate at which polygons enter the band. This includes the shrinking of larger polygons, which enter the band at rate *v*(*δA*)*P* (*δA*) and the creation of new polygons due to splitting of polygons with area between (0, 2*δA*):

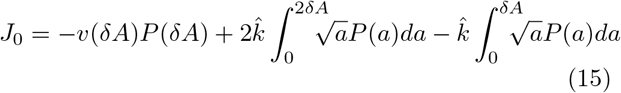

Here, the second term corresponds to 2 new polygons created by splitting and the last term accounts for the fact that the original polygon vanishes when a splitting event occurs. If we assume that the distribution function for small areas is a power law: *P* (*A*) = *A*^*μ*^, then the above equation can be expressed as:

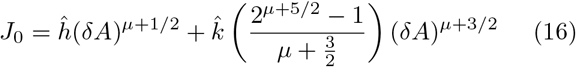

When very small areas are being considered (*δA →* 0), the first term is dominant, so we have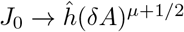. In order to have a finite flux of disappearing polygons, countering the production of new polygons throughout the system, we must set *μ* = *−* 1*/*2.

Therefore the distribution function for polygons of small area can be written as

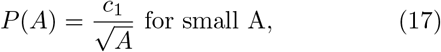

with *c*_1_ a constant. Converting to a logarithmic distribution, 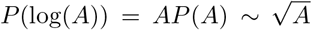. This square root scaling matches the distribution of small polygon areas measured for simulations of liquid networks (Fig. 3A and Fig. 4H).

#### Large area scaling

For very large values of the area *A*, the polygon dynamics are described by a balance between growth and splitting. We neglect the last term in Eq. 14, assuming that *P* (2*A*) *≪ P* (*A*) so that the production of very large polygons by splitting events is rare. This assumption is self-consistent with an exponentially decaying distribution for large areas. Plugging the growth velocity from Eq. 13a into the truncated form of Eq. 14 gives the steady-state distribution of polygon areas as:

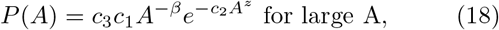

where 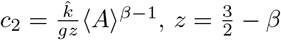, and *c*_3_, *c*_1_ constants.

#### Full distribution by stitching together the two limits

In order to construct the complete distribution, the two limits for small and large area are married together at some intermediate area *A*^*\**^. Enforcing continuity of the distribution requires 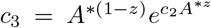and gives the following functional form for the full distribution:

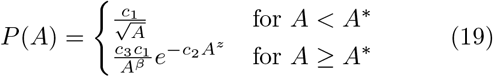

The remaining unknowns are *c*_1_, *A*^*\**^, and ⟨*A*⟩ (which also appears in *c*_2_). In order to estimate *A*^*\**^ and ⟨*A*⟩, we first note that the total number of polygons must remain fixed at steady state. Thus, the flux of vanishing small polygons must equal the flux of newly created polygons via splitting: *J*_0_ = *J*_split_. The flux of vanishing polygons is:

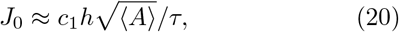

and the flux of newly created polygons due to splitting is:

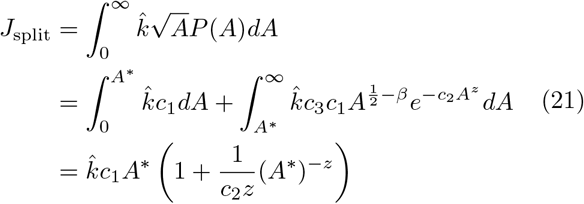

Additionally, an expression for the average area is obtained via 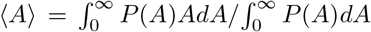. Through this matching of fluxes and the definition of average area, we obtain two equations:

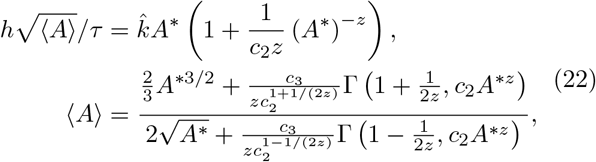

where Γ is the incomplete Gamma function. These can be solved numerically for *A*^*\**^ and ⟨*A*⟩ using parameters *g, h*, and *β* from the effective growth law. Finally, the prefactor *c*_1_ is set by normalizing the full distribution 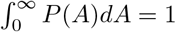.

Using the non-dimensionalized growth law extracted from liquid network simulations (Fig. 4E, Eq. 4), this gives *A*^*\**^ *≈* 0.12𝓁^2^ and ⟨*A*⟩ ≈ 0.23𝓁^2^. We note that the predicted average area is similar to the actual value of ⟨*A*⟩ ≈ 0.29𝓁^2^ obtained for simulated liquid networks (Fig. 2D).

The area *A*^*\**^ corresponds to the peak in the area distribution function *P* (*A*) (see Fig. 4H) and thus gives an estimate of the most ‘typical’ polygon area. For COS7 ER, we have the length scale 𝓁 *≈* 1.95 *μ*m, giving the predicted typical area *A*^*\**^ ≈ 0.46 *μ*m^2^ in the liquid network model. For comparison, the peak in the measured area distribution for ER polygons in COS7 cells is 0.72 *±* 0.32 *μ*m^2^ (mean and standard deviation across cells).

### Supplemental Video Captions

- **Video 1:** Confocal movie of COS7 cell expressing fluorescent endoplasmic reticulum marker (KDEL_mcherry) demonstrating junction sliding and ring closure.
- **Video 2:** Four simulated liquid networks each with a different combination of junction mobility and tubule spawning rate leading to varying tubule density and rearrangement timescales. *b* = 10^*−*3^ *μ*m/s and 10^*−*2^ *μ*m/s in top and bottom row; *k* = 5 *×* 10^*−*4^ *μ*m^*−*1^s^*−*1^ and 5 *×* 10^*−*3^ *μ*m^*−*1^s^*−*1^ in left and right column, respectively.
- **Video 3:** Time evolution of polygon areas in a liquid network, highlighting growth and shrinking dynamics of the polygon population. Each circle represents the extracted area of a polygon centered at that location. Small circles shrink and large circles grow until they are split.
- **Video 4:** Liquid network simulation with semi-persistent pinning to static structures. Here, *b* = 0.02 *μ*m/s, *k* = 0.005 *μ*m^*−*1^s^*−*1^, *k*_*p*_ = 10 s^*−*1^, *k*_*u*_ = 0.01 s^*−*1^, leading to an average of 1000 pins at steady-state.
- **Video 5:** Example 6.3 *×* 6.3 *μ*m region of peripheral ER, visualized for 40 s in order to count new tubule spawning events and calculate tubule spawning rate. Region is extracted from a confocal time-lapse image of COS7 cell expressing fluorescent endoplasmic reticulum marker (KDEL_mcherry). Video is shown in 2*×* real-time.

